# Durability of SARS-CoV-2-specific T cell responses at 12-months post-infection

**DOI:** 10.1101/2021.08.11.455984

**Authors:** Zhongyan Lu, Eric D. Laing, Jarina Pena-Damata, Katherine Pohida, Marana S. Tso, Emily C. Samuels, Nusrat J. Epsi, Batsukh Dorjbal, Camille Lake, Stephanie A. Richard, Ryan C. Maves, David A. Lindholm, Julia Rozman, Caroline English, Nikhil Huprikar, Katrin Mende, Rhonda E. Colombo, Christopher J. Colombo, Christopher C. Broder, Anuradha Ganesan, Charlotte A. Lanteri, Brian K. Agan, David Tribble, Mark P. Simons, Clifton L. Dalgard, Paul W. Blair, Josh Chenoweth, Simon D. Pollett, Andrew L. Snow, Timothy H. Burgess, Allison M.W. Malloy, the EPICC COVID-19 Cohort Study Group

**Affiliations:** Department of Pediatrics, Uniformed Services University of the Health Sciences, Bethesda, MD, USA, 20814; Henry M. Jackson Foundation for the Advancement of Military Medicine, Inc., Bethesda, MD, USA, 20817; Department of Microbiology and Immunology, Uniformed Services University of the Health Sciences, Bethesda, MD, USA, 20814; Department of Pharmacology & Molecular Therapeutics, Uniformed Services University of the Health Sciences, Bethesda, MD, USA, 20814; Infectious Disease Clinical Research Program, Department of Preventive Medicine and Biostatistics, Uniformed Services University of the Health Sciences, Bethesda, MD, USA, 20814; Department of Anatomy, Physiology & Genetics, Uniformed Services University of the Health Sciences, Bethesda, MD, USA, 20814; Austere Environments Consortium for Enhanced Sepsis Outcomes, Henry M. Jackson Foundation, Bethesda, MD, USA, 20817; Naval Medical Center San Diego, San Diego, CA, USA, 92134; Brooke Army Medical Center, JBSA Fort Sam Houston, TX, USA, 78234; Madigan Army Medical Center, Tacoma, WA, USA, 98431; Walter Reed National Military Medical Center, Bethesda, MD, USA, 20814

**Keywords:** COVID-19, SARS-CoV-2, 12-months, T cell, antibody, memory, cytotoxicity, polyfunctionality, durability

## Abstract

**Background:** Characterizing the longevity and quality of cellular immune responses to SARS-CoV-2 is critical to understanding immunologic approaches to protection against COVID-19. Prior studies suggest SARS-CoV-2-specific T cells are present in peripheral blood 10 months after infection. Further analysis of the function, durability, and diversity of the cellular response long after natural infection, over a wider range of ages and disease phenotypes, is needed to further identify preventative and therapeutic interventions.

**Methods:** We identified participants in our multi-site longitudinal, prospective cohort study 12-months post SARS-CoV-2 infection representing a range of disease severity. We investigated the function, phenotypes, and frequency of T cells specific for SARS-CoV-2 using intracellular cytokine staining and spectral flow cytometry. In parallel, the magnitude of SARS-CoV-2-specific antibodies was compared.

**Results:** SARS-CoV-2-specific antibodies and T cells were detected at 12-months post-infection. Severity of acute illness was associated with higher frequencies of SARS-CoV-2-specific CD4 T cells and antibodies at 12-months. In contrast, polyfunctional and cytotoxic T cells responsive to SARS-CoV-2 were identified in participants over a wide spectrum of disease severity.

**Conclusions:** Our data show that SARS-CoV-2 infection induces polyfunctional memory T cells detectable at 12-months post-infection, with higher frequency noted in those who originally experienced severe disease.

## Introduction

Understanding the development and durability of protective immune responses against SARS-CoV-2 remains critical as we seek global reduction in disease burden. Antibody responses induced during a primary SARS-CoV-2 infection have been shown to wane, but may be present in the circulation up to twelve months post symptoms onset (PSO) [1, 2]. Thus far, SARS-CoV-2- specific T cells have been detected up to 10 months PSO [3, 4]. Intriguingly, SARS-CoV-1-specific T cells have been identified 17 years post-infection, suggesting the potential for very long-lived T cell memory [5]. Antibodies have been effective in reducing the disease burden of SARS-CoV-2, however, infection still occurs requiring the recruitment of T cells to clear infected cells. In order to utilize the specificity and potent viral clearance of T cells, further knowledge is required regarding viral antigen specificity, memory differentiation and longevity [6].

Studies on SARS-CoV-2-specific T cells within two months PSO suggests that potent antigen-specific T cells are associated with mild disease, whereas a lack of these antiviral cells or a delay in development is associated with severe disease [7, 8]. Peng et al. [9] showed that individuals who had mild disease had a higher ratio of polyfunctional CD8 T cells compared to CD4 T cells around 42 days PSO, suggesting that potent SARS-CoV-2-specific CD8 T cells may be protective. Analysis of functional T cell responses at memory time points is needed to provide insight into their cytolytic potential and role in protection upon re-infection that prior studies relying on activation-induced markers (AIM) [6] could not.

The T cell response to SARS-CoV-2 has been shown to recognize epitopes across multiple viral proteins including the spike glycoprotein (S), nucleocapsid (N), membrane (M), and small envelope (E) proteins, as well as other nonstructural proteins (nsp) [10, 11]. Due to the ability of T cells to recognize structural and nonstructural proteins that are less susceptible to antibody-dependent mutational pressure, T cells may provide cross-reactive protection against other coronaviruses as well as SARS-CoV-2 variants [12, 13].

In this study, we analyzed the durability and functional characteristics of the SARS-CoV-2-specific memory T cell response at 12-months PSO derived from a prospective, longitudinal cohort of United States Military Health System (MHS) beneficiaries, including active-duty military and dependents with varying severity of disease. Understanding the durability and functional response of SARS-CoV-2-specific T cells can help determine how the humoral and cellular components of antiviral immunity work synergistically to prevent future infection and inform vaccine strategies.

## Methods and Materials

### Study Participants

Individuals were enrolled into the Epidemiology, Immunology, and Clinical Characteristics of Emerging Infectious Diseases with Pandemic Potential (EPICC) study, a prospective, longitudinal cohort study, if they were exposed to, had symptoms consistent with, or had documented SARS-CoV-2 infection beginning in March of 2020. Samples from four of ten military treatment facilities (MTFs), which included Walter Reed National Military Medical Center (Bethesda, MD), Brooke Army Medical Center (San Antonio, TX), Naval Medical Center San Diego (San Diego, CA), and Madigan Army Medical Center (Tacoma, WA) were included based on peripheral blood mononuclear cells (PBMCs) obtained 12-months PSO and absence of evidence for re-infection. Infection was confirmed by reverse-transcriptase polymerase chain reaction (RT-PCR) and serologic response. Study participants included in this analysis were followed for 12-months PSO with biological samples and clinical questionnaire data collected throughout the duration of the study. Receipt of a SARS-CoV-2 vaccine was identified by report and confirmed by medical data repository review. Study participants who received a SARS-CoV-2 vaccine prior to the 12-month peripheral blood collection were excluded from S-specific T cell and antibody analysis. Study participants were evaluated for re-infection by documented PCR positive for SARS-CoV-2 or a significant rise in both S and N-specific antibody responses. Participants who were re-infected were excluded for comparative T cell and antibody analysis in the current study. The protocol was approved by the Uniformed Services University Institutional Review Board (IDCRP-085), and all subjects or their legally authorized representative provided informed consent to participate.

### PBMC and serum preparation

PBMCs were isolated from peripheral whole blood collected in acid citrate dextrose (ACD) tubes at 12-months PSO. PBMCs were purified using a Ficoll-Histopaque (Fisher Scientific, NH) gradient. Cells were preserved in 90% fetal bovine serum (Sigma-Aldrich, MO) and 10% dimethyl sulfoxide (DMSO, Sigma-Aldrich, MO) and stored in liquid nitrogen. Serum was isolated from blood collection in EDTA tubes.

### Measurement of antigen presenting cells

One million thawed PBMCs were stained in PBS at 4°C for 20 minutes for antigen presenting cells (APCs) using the antibodies listed in Supplemental Table 1 and were acquired on a CYTEK Aurora 5-laser spectral flow cytometer (CYTEK Biosciences, CA).

### Antigen-specific T cell analysis

For identification of antigen-specific T cells, isolated PBMCs were cultured in 96-well plates and stimulated overnight with peptide pools derived from selected viral proteins. Peptide pools were comprised of 15-mer peptides overlapping by 11 amino acid residues covering the S, M, N, and E proteins of SARS-CoV-2 (JPT, Germany), the S protein of endemic human coronavirus (hCoV) strains: HKU1, 229E, NL63 and OC43; and a CMV peptide pool including pp50, pp65, IE1, IE2 and envelope glycoprotein B (Mabtech, OH) at a final concentration of 1 mg/ml for each individual peptide (Supplemental Table 2). Monensin (BD Biosciences, CA; BioLegend, CA) was added to the wells 1 hour after peptide addition to prevent cytokine secretion as recommended by the manufacturer. Ten minutes after addition of the peptide pools, CD107a antibody (BioLegend, CA) was added. Cell surface markers were identified with the antibodies listed in Supplemental Table 3. Surface staining was followed by cell permeabilization using FoxP3/transcription factor staining buffer set (Thermo Fisher Scientific, MA) at 4°C for at least 30 minutes. Antibodies used for intracellular cytokine staining (ICS) (Supplemental Table 3) were then added. PBMCs cultured in medium without peptide stimulation served as the negative control, or with CytoStim (Miltenyi Biotec, CA) as the positive control for the assay. Samples were acquired on a CYTEK Aurora (CYTEK Biosciences, CA). All cytometric data were analyzed using FlowJo software (BD Biosciences, CA). Samples were considered positive if the frequency of IFNγ+ T cells was 2-fold higher than the medium control and greater than 0.01% of CD4 or CD8 T cells after subtracting the medium control value [14].

### Multiplex microsphere-based immunoassay screening procedures

Detailed experimental procedures of SARS-CoV-2 S and N protein-based multiplex microsphere immunoassays have been previously described [2, 15]. Briefly, diluted serum and capillary blood samples were added to 96-well microtiter plates containing antigen-coupled microspheres and tested in technical duplicates. After 45 minutes of agitation, wells were washed, and biotin-conjugated goat anti-human IgG (Thermo Fisher Scientific, Waltham, MA) diluted in PBS + 0.05% Tween20 (PBST) was added to each well. Wells were subsequently washed again, then streptavidin-phycoerythrin (1:1000 in PBST) (Bio-Rad, Hercules, CA) was added to each well. A final wash was performed after 45 minutes of incubation, and antigen-antibody complexes were analyzed on BioPlex 200 multiplexing systems (Bio-Rad) for IgG binding, and median fluorescence intensity (MFI) values were reported for specificity to SARS-CoV-2 S and N protein. Participants were excluded from 12-months PSO evaluation based on observed increases in SARS-CoV-2 S protein binding from 6-12 months collections across longitudinal serum samples and an accompanying rise in N protein IgG results, suggesting possible re-infections.

### Statistical analysis

Data were analyzed using GraphPad Prism 9 (GraphPad Software Inc, CA). Two-group test significance levels were calculated using Mann-Whitney analysis. Correlation coefficients and significance levels were calculated using Spearman rank correlation. A p-value < 0.05 was considered statistically significant.

## Results

### Study cohort

From our cohort, we identified 29 patients who had COVID-19 approximately 1 year prior, and did not have evidence of re-infection prior to a peripheral blood collection at 12-months PSO. Disease severity was classified by inpatient or outpatient status during the acute phase of illness. The age range of the study participants was 20-72 years of age and the overall distribution of age did not vary significantly (p = 0.1) between inpatients (median age 50.6 with a range of 21.4 to 72.4) and outpatients (median age 44.6 with a range of 20.1 to 60.1), nor did sex (Table 1, Supplemental Table 4). Inpatients, however, were more likely to have a comorbidity with respiratory and pulmonary illnesses being the most common. 6/14 study inpatients received intermittent intranasal or inhaled corticosteroids for underlying conditions, in contrast to 1/15 outpatient. Two study participant received systemic steroids during acute illness (Table 1, Supplemental Table 4). Data on S-specific humoral and cellular responses was not included for individuals receiving vaccination prior to the 12-month blood drawn. Patients hospitalized during acute illness were further dichotomized into those who required supplemental oxygen during hospitalization and those who did not. Further demographic and clinical information are provided in Supplemental Table 4.

**Table 1.** Demographic table of participants

|  | Hospitalized (N=14) | Outpatient (N=15) | Total (N=29) |
| --- | --- | --- | --- |
| **Male** | 8 (57.1%) | 8 (53.3%) | 16 (55.2%) |
| **Race** |  |  |  |
| Asian | 2 (14.3%) | 2 (13.3%) | 4 (13.8%) |
| Black | 8 (57.1%) | 1 (6.7%) | 9 (31.0%) |
| Hispanic | 1 (7.1%) | 5 (33.3%) | 6 (20.7%) |
| Other | 0 (0.0%) | 1 (6.7%) | 1 (3.4%) |
| White | 3 (21.4%) | 6 (40.0%) | 9 (31.0%) |
| **Age** |  |  |  |
| Median (Q1, Q3) | 50.6 (45.3, 55.8) | 44.6 (39.7, 50.9) | 46.4 (40.1, 53.5) |
| Min - Max | 21.4 - 72.4 | 20.1 - 60.2 | 20.1 - 72.4 |
| **Age group** |  |  |  |
| <18 | 0 (0.0%) | 0 (0.0%) | 0 (0.0%) |
| 18-44 | 3 (21.4%) | 8 (53.3%) | 11 (37.9%) |
| 45-64 | 10 (71.4%) | 7 (46.7%) | 17 (58.6%) |
| 65+ | 1 (7.1%) | 0 (0.0%) | 1 (3.4%) |
| **Military affiliation** |  |  |  |
| Active duty | 2 (14.3%) | 5 (33.3%) | 7 (24.1%) |
| Dependent | 6 (42.9%) | 6 (40.0%) | 12 (41.4%) |
| Retired military | 6 (42.9%) | 4 (26.7%) | 10 (34.5%) |
| **Any comorbidities** | 13 (92.9%) | 7 (46.7%) | 20 (69.0%) |
| **Multiple comorbidities** | 4 (28.6%) | 2 (13.3%) | 6 (20.7%) |
| **Charlson comorbidity index** |  |  |  |
| Median (Q1, Q3) | 1.0 (0.0, 1.5) | 0.0 (0.0, 1.5) | 1.0 (0.0, 1.75) |
| Min - Max | 0 - 6 | 0 - 3 | 0 - 6 |

### Convalescent T cell and antibody responses are detectable 12-months post symptoms onset

Identification of SARS-CoV-2-specific T cells was performed by stimulating PBMCs with peptide pools derived from the S, N, M, and E proteins. Stimulation of PBMCs with these peptide pools demonstrated the presence of SARS-CoV-2-specific CD4 and CD8 T cells at 12-months PSO by ICS (Figure 1A; Supplemental Figure 1A) in 22/29 (75.9%) of study participants. Only three of 29 participants had a measurable response to E (data not shown); therefore, the presented data focused on S, M, and N. Stimulation with the CMV peptide pool demonstrated CMV-specific CD4 and CD8 T cell responses in 73.9% of study participants. Overall, SARS-CoV-2-specific CD4 T cells were more frequently identified in the peripheral blood compared to SARS-CoV-2-specific CD8 T cells. In addition, T cell responses to endemic hCoV strains; HKU1, 229E, OC43, and NL63 were sporadic, with no differential distribution or magnitude difference between inpatients and outpatients identified (Table 2).

**Figure 1.**
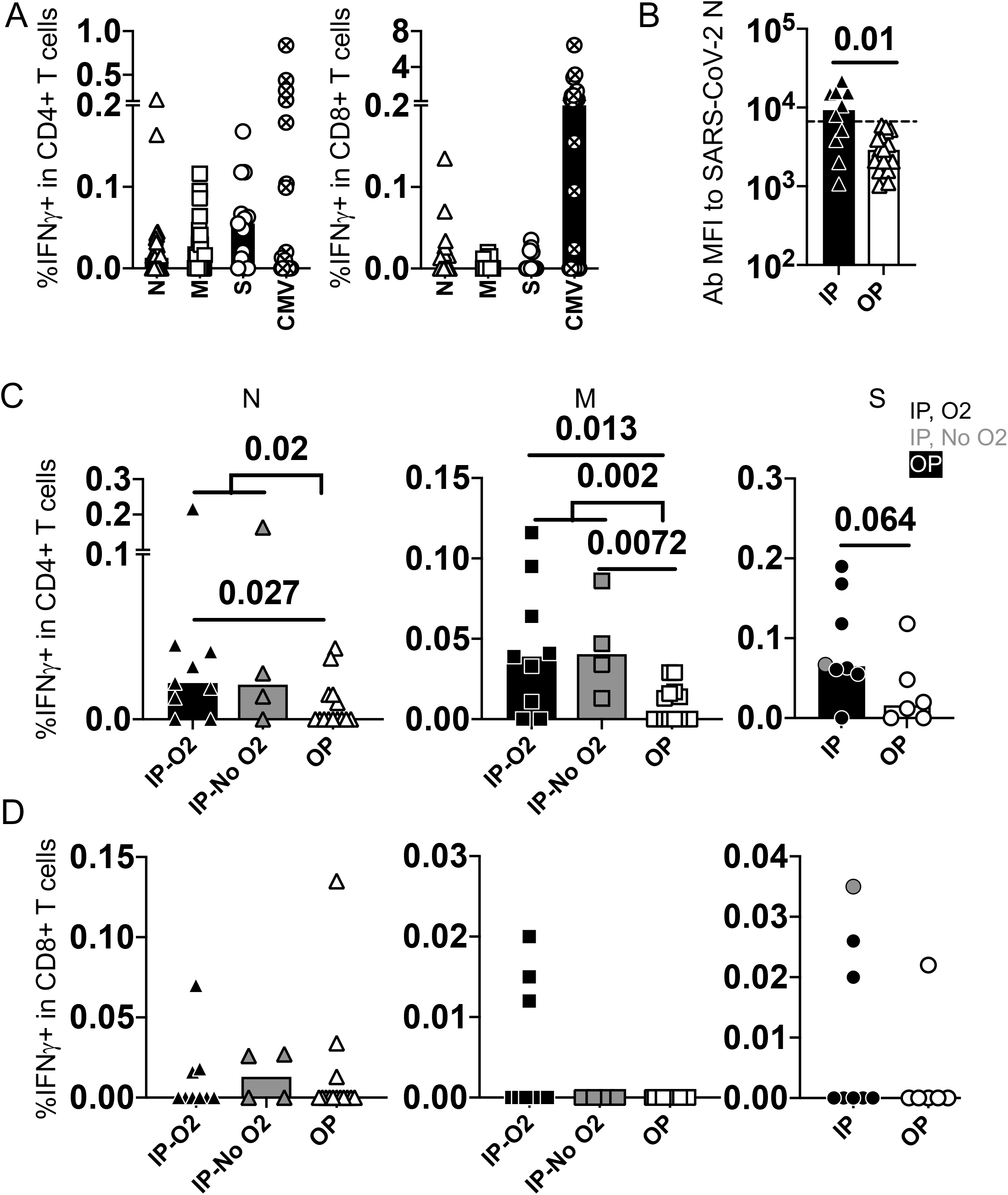
Adaptive immune responses to SARS-CoV-2 viral components at 12-months post symptoms onset. **A**. SARS-CoV-2-specific CD4 and CD8 T cell responses were detected at 12- months PSO by expression of IFNγ upon peptide stimulation and intracellular cytokine staining. **B**. Antibodies to SARS-CoV-2 nucleocapsid protein at 12-months post-infection. Threshold shows the global MFI of antibodies to endemic hCoVs using the same sera in the same assay. **C**. Frequency of CD4 T cell responses to the tested viral components N, M, and S in all patient groups. **D**. Frequency of CD8 T cell responses to the tested viral components N, M, and S in all patient groups. Bars represent the median value in each group. P values in the plots indicate the significance by Mann-Whitney test.

**Table 2.** Frequencies of T cell responses to endemic CoV d SARS-CoV-2 viral components

|  | HKU-S | NL63-S | OC43-S | 229E-S |  | SARS-CoV-2 N | SARS-CoV-2 M | SARS-CoV-2 E |
| --- | --- | --- | --- | --- | --- | --- | --- | --- |
| Inpatient, O2 | 3 of 7 | 4 of 6 | 3 of 6 | 2 of 7 |  | 7 of 11 | 9 of 11 | 0 of 11 |
| Inpatient, no O2 | 3 of 6 | 3 of 6 | 3 of 6 | 2 of 6 |  | 3 of 3 | 3 of 3 | 2 of 3 |
| outpatient | 4 of 14 | 4 of 14 | 5 of 14 | 4 of 14 |  | 6 of 15 | 7 of 15 | 1 of 15 |

The SARS-CoV-2-specific antibody response against S and N proteins were also present in both unvaccinated inpatient and outpatient groups at 12-months PSO (Figure 1B, Supplemental Figure 1B). We found that the antibody response against N in inpatients was higher than in outpatients, in whom it waned to levels below the threshold of endemic hCoV N responses (P = 0.01). These findings demonstrate that the humoral and cellular responses to SARS-CoV-2 exhibit durability at 12-months PSO, but vary by severity of disease and antigen specificity.

Stratification of study participants by inpatient and outpatient status at peak disease severity demonstrated that individuals with more severe disease exhibited the highest frequency of CD4 T cells responsive to N (P = 0.02), M (P = 0.002), and S (P = 0.064) proteins, regardless of oxygen requirement (Figure 1C). Of the inpatients, 11 of 13 (84.6%) exhibited T cell responses specific for either N or M (P30 and P41 did not exhibit T cell responses above assay cutoff). In contrast, 7 of 14 (50.0%) outpatients exhibited CD4 T cell responses to N or M. Individuals who exhibited T cell specificity for either N or M at 12-months were also more likely (P = 0.0007) to simultaneously recognize other SARS-CoV-2 epitopes in N or M (Supplemental Figure 1C). Compared with the CD4 T cell response, the CD8 T cell response, as measured by IFNγ production, was lower in magnitude and did not correlate with severity of disease (Figure 2D).

**Figure 2.**
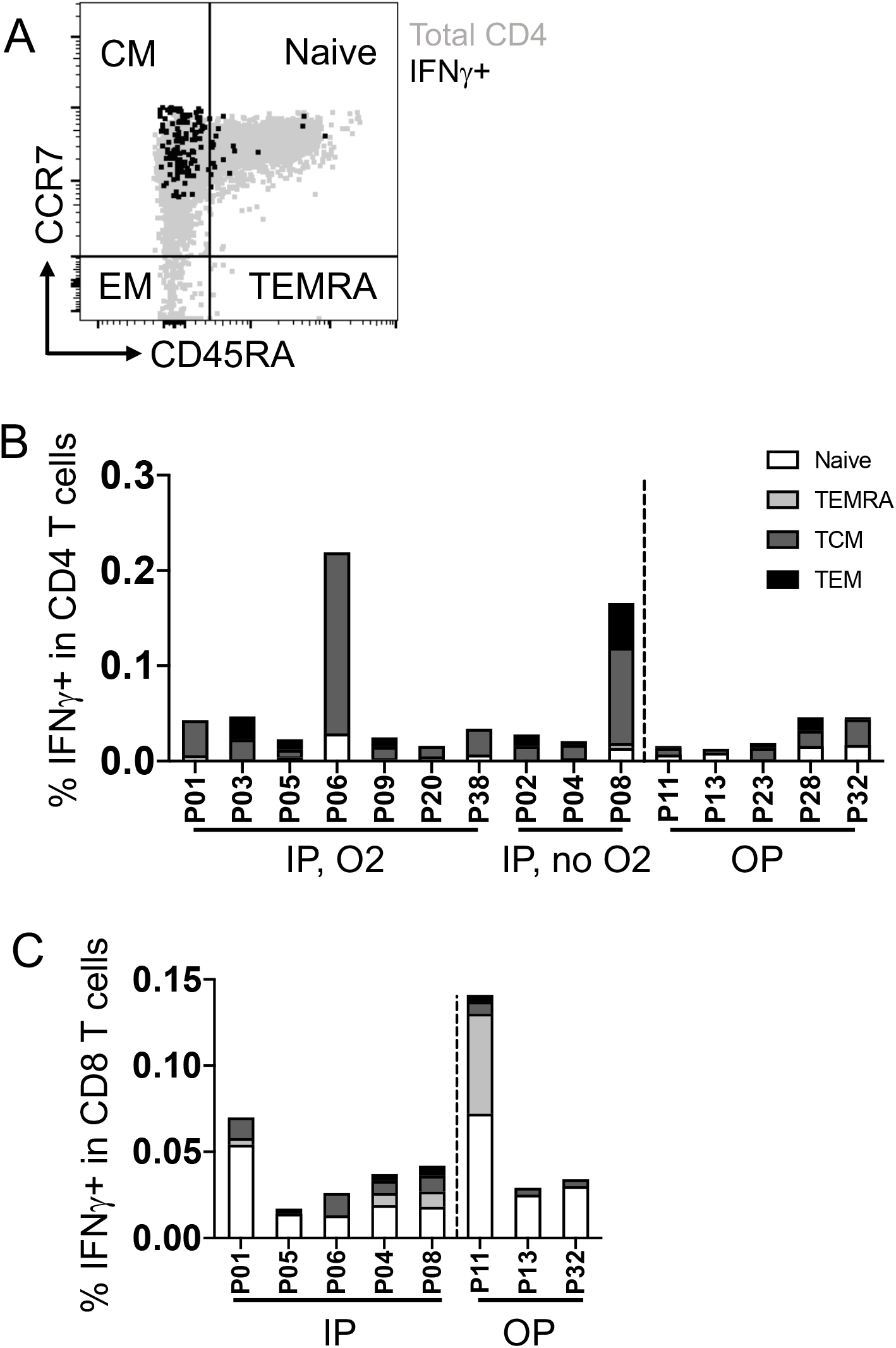
Memory phenotype characterization of SARS-CoV-2-specific T cell responses. **A**. A representative flow plot of the essential markers used for memory phenotyping. **B**. Memory phenotypes of IFNγ+ CD4 T cells to SARS-CoV-2 N protein as an example in each patient separated by patient group. **C**. Memory phenotypes of IFNγ+ CD8 T cells to SARS-CoV-2 N protein in each patient separated by patient group.

Quantification of the APCs and frequency of total T cells at 12-months PSO were measured by flow cytometry (Supplemental Figure 1D) to determine whether differences exhibited between study groups were influenced by APC or T cell availability. Our data show that the ratio of total CD4 and CD8 T cells (Supplemental Figure 1E), and the frequency of classical APCs, including conventional dendritic cells 1 and 2 (cDC1/2), plasmacytoid DCs (pDCs) and monocytes (Supplemental Figure 1F), were similar among patient groups at 12-months PSO and healthy controls.

### SARS-CoV-2-specific T cells at 12-months post symptoms onset predominantly exhibit central memory characteristics

SARS-CoV-2-specific T cells identified by IFNγ production in response to SARS-CoV-2 peptides were characterized with established phenotypic markers of differentiation as previously described [6]. Expression of the lymph node homing receptor, CCR7, coupled with the absence of CD45RA expression is characteristic of central memory T cells [16] (Figure 2A), which are also CD27+CD28+ [17]. This was the predominant phenotype exhibited by CD4 T cells responding to N (Figure 2B; Supplemental Figure 2A) and was distinct from the predominance of naïve T cells in the total CD4 T cell population. The phenotypic distribution of CD4 T cells responding to M and S was similarly skewed towards a central memory phenotype (Supplemental Figure 2B). In contrast, CD8 T cells specific for N and S derived from multiple compartments, including naïve (CCR7+CD45RA+), effector memory T cells re-expressing CD45RA (TEMRA), central and effector memory phenotypes (Figure 2C; Supplemental Figure 2C).

Together these findings demonstrate that the SARS-CoV-2-specific T cell responses exhibit a longevity of at least 12-months, with a predominance of central memory CD4 T cells specific for S, N and M proteins. SARS-CoV-2-specific CD8 T cells were less frequent in the peripheral blood and exhibited more diverse memory phenotypes. For both CD4 and CD8 T cells, the severity of disease during acute illness was not reflected in distinct memory phenotype differentiation at 12-months PSO.

### Cytokine profiles indicate that SARS-CoV-2-specific memory T cells are polyfunctional

T cells were evaluated for the expression of multiple cytokines in response to the distinct SARS-CoV-2 peptide pools to determine their functional potential. CD4 T cells responsive to N exhibited expression of both IFNγ and IL2, indicating a polyfunctional response in all study participants in both inpatient and outpatient severity groups at 12-months PSO (Figure 3A). Overall, inpatients had a higher frequency of cytokine-expressing CD4 T cells, but the frequency of polyfunctional CD4 T cells did not differ significantly between severity groups (Figure 3A). CD4 T cell responses against M and S exhibited similar frequencies of polyfunctional T cells, suggesting that epitope specificity did not predict polyfunctionality (Supplementary Figure 3A-B). At 12-months, we did not detect expression of IL17A, IL21, IL4 or IL13 by CD4 T cells in response to the evaluated peptide pools (data not shown) based on the assay cut-off, suggesting that type 1 helper (Th1) CD4 T cells were dominant over Th2, Th17, or circulating T follicular helper cells (cTfh).

**Figure 3.**
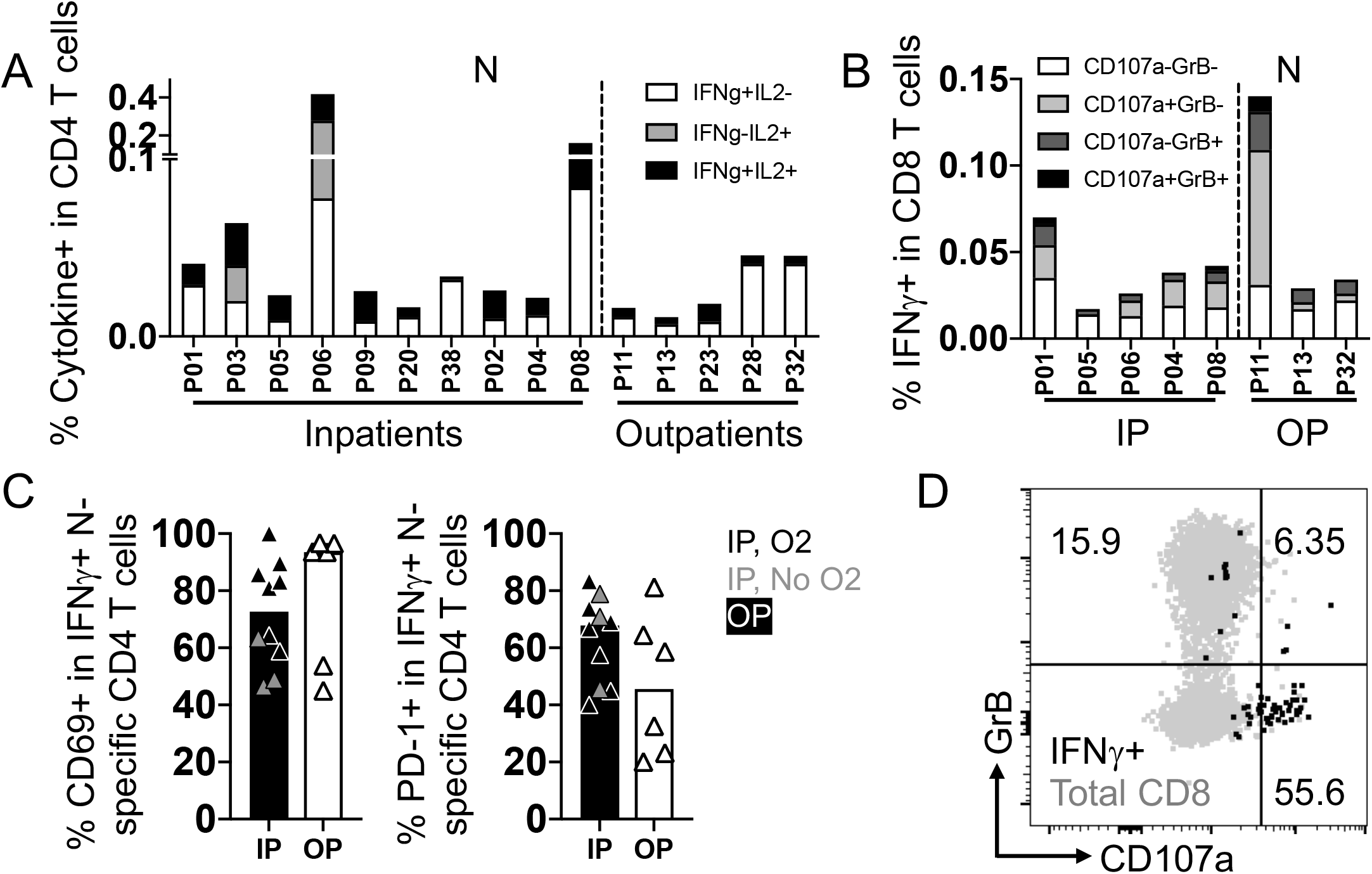
Polyfunctionality, cytotoxicity, and activation of SARS-CoV-2-specific CD4 or CD8 T cells. **A**. Polyfunctionality of CD4 T cell responses to SARS-CoV-2 N protein measured by IL2 and IFNγ. **B**. Cytotoxicity of IFNγ+ CD8 T cells to SARS-CoV-2 N protein measured by expression of intracellular Granzyme B and membrane CD107a. **C**. Frequencies of CD69 and PD-1 in SARS-CoV-2 N-specific IFNγ+ CD4 T cells in each patient group. Grey dots in the inpatient group represent inpatients without oxygen supplementation. **D**. A representative flow plot of cytotoxic CD8 T cells by the expression of Granzyme B and CD107a.

CD8 T cells expressing IFNγ were further evaluated for cytolytic potential by the expression of Granzyme B and surface exposure of the lysosome-associated membrane protein CD107a, a marker of degranulation (Figure 3D). The CD8 T cell response to N and S displayed heterogeneity in cytolytic potential, with cells generally expressing either marker in response to their cognate antigens (Figure 3B; Supplemental Figure 3B). Again, we saw no difference in frequency and function of the SARS-CoV-2-specific CD8 T cells based on disease severity.

We further looked into the activation features of the IFNγ+ CD4 T cells. Upon peptide stimulation, SARS-CoV-2-specific IFNγ+ CD4 T cells upregulated CD69 and PD-1 compared to the total CD4 T cell population, and was similar in inpatients and outpatients (Figure 3C; Supplementary Figure 3C-D). The other activation markers, including CD38, HLA-DR, or CTLA-4, in the IFNγ+ T cells are comparable to the total CD4 T cell population (Supplementary Figure 3C).

Together these data suggest that the memory CD4 T cell response against SARS-CoV-2 predominantly exhibits a Th1 type profile, and that SARS-CoV-2-specific memory CD8 T cells display cytolytic potential at 12-months.

## Discussion

Here we show that SARS-CoV-2-specific T cell responses are detected at 12-months PSO, extended our prior understanding of the durability of the T cell response against SARS-CoV-2 [4]. The frequency of SARS-CoV-2-specific CD4, but not CD8, T cells was higher at 12-months PSO in individuals who experienced severe disease compared to those who had mild disease at the acute phase. Importantly, memory SARS-CoV-2-specific CD4 T and CD8 T cells exhibited polyfunctionality and cytotoxicity, respectively, suggesting strong recall responses. Our findings and that of others also show that antibodies to SARS-CoV-2 persist 12-months PSO, but wane rapidly [1, 2, 18] and in the case of N-specific antibodies may be undetectable.

Overarchingly, we identified SARS-CoV-2-specific T cell responses in 75.9% of study participants at 12-months PSO using peptide pools derived from SARS-CoV-2 structural proteins N, M, E, and S. Other studies have shown that epitopes in SARS-CoV-2 N, M, and S, together with nsp3, 4, ORF3a are recognized by CD4 and CD8 T cells representing the majority of T cell responses at convalescence, while SARS-CoV-2 E was less frequently recognized [11, 19, 20]. Although we did not map T cell epitopes, we overall observed the same trend of T cell responses to corresponding peptide pools used in this study, but at a much later time point. As the magnitude of CD8 T cell responses contracts within a month of disease onset in humans vaccinated with live yellow fever virus and smallpox vaccines [21, 22], similarly in COVID-19 patients [23], SARS-CoV-2-specific CD8 T cells in the peripheral blood are of low frequency at one year PSO. Importantly, T cell recognition of multiple epitopes was common within study participants, suggesting broad epitope recognition.

Study participants who were hospitalized during acute infection demonstrated the highest frequency of SARS-CoV-2-specific CD4 T cell responses and antibodies to SARS-CoV-2 N. Why severe COVID-19 is associated with a higher memory response for CD4 T cells remains unclear. Since severe disease is associated with higher viral burden [24-26], one hypothesis is that higher viral load at acute infection increases stimulation and antigen availability for T cells expanding more durable responses. However, due to differences in timing of nasal swab acquisition and the size of this subcohort, we were unable to correlate T cell longevity and viral load at acute infection. Published data have yet to show that initial SARS-CoV-2 viral load correlates to the longevity of cellular immunity. Indeed, the longevity of memory T cells is affected by multiple factors including the cytokines such as IL7 and IL15 (reviewed in [27]) as well as the density of antigens presented by APCs, including B cells [28].

Importantly, our data show that SARS-CoV-2-specific CD4 T cells present in individuals infected 12-months prior exhibit differentiation toward central memory phenotypes, activation by expression of CD69 and PD-1, and polyfunctionality by expression of IFNγ and IL2. In other viral infections such as influenza, polyfunctional CD4 T cells expressing IFNγ, IL2, and TNFα, are more frequent in convalescent patients who better controlled infection [29]. COVID-19 patients with milder symptoms also had more polyfunctional T cells expressing IFNγ, IL2, and TNFα [30] while severe and critical patients tended to have restricted functional T cells 1-2 months PSO [31]. CD4 T cells 1-2 months after SARS-CoV-2 infection also upregulated CD69, CD38, HLA-II, CTLA-4, and PD-1, as an indication of activation [31, 32]. While the polyfunctionality and activation of long-term memory CD4 T cells is less well characterized in other viral infections and could be impacted by factors such as age and antigen concentration [33-35], in our study, the SARS-CoV-2-specific CD4 T cells between severity study groups were similarly polyfunctional and equally expressed CD69 and PD-1 at 12-months PSO. In addition, although CD8 T cells were less frequent compared to CD4 T cells at 12-months PSO, SARS-CoV-2-specific CD8 T cells exhibited cytotoxicity while CD4 T cells did not (data not shown). Memory CD4 T cells are, however, important in promoting CD8 T cell cytotoxicity and viral clearance (reviewed in [36]).

SARS-CoV-2 shares epitopes with endemic hCoVs including HKU1, 229E, NL63, and OC43 [32, 37]. Our analysis was restricted to hCoV peptide pools comprised of S from these viruses and we detected low frequencies of T cell recognition of endemic hCoV that did not differ between patient severity groups. Analysis of additional structural and non-structural protein-derived peptides in larger study cohorts may yield more information regarding the correlation of cross-reactive T cell responses. Moreover, T cells can provide ostensibly broad coverage over SARS-CoV-2 variants of concern, whereas antibodies primarily target S [13]. Data has shown that associated mutations can develop and emerge in the receptor binding domain and other sites of the S glycoprotein that limit prior antibody binding [38-40]. However, due to the unique requirements of peptide presentation in the context of an HLA molecule, distinct epitope-HLA haplotype interactions between individuals present an overwhelming number of mutational target sites for which the virus is unable to evade.

Limitations to our study include the challenge of obtaining biological samples at 12-months PSO prior to vaccination in our MHS cohort, which had timely and reliable access to vaccines. Fifty percent of individuals in our cohort were vaccinated prior to a 12-month draw (Table 1). For these individuals we did not include their S-specific T cell or antibody responses, but did include their N, M and E humoral or cellular responses since these are not components of the mRNA-based vaccines (the vaccines primarily available to our study population). A strength of our cohort is that study participants are followed within the MHS and their epidemiologic, clinical and COVID-19 testing records, and vaccination status are maintained primarily from electronic health record. In addition, sera were analyzed longitudinally to avoid re-infected participants for 12-months T cell analysis.

In summary, our data show that both SARS-CoV-2 humoral and cellular responses are measurable 12-months PSO in individuals across a spectrum of disease phenotypes. The magnitude of the CD4 T cell and antibody responses more closely correlated with acute disease severity. Importantly, the memory phenotype and polyfunctional response of the SARS-CoV-2 specific T cells did not differ by disease severity. The breadth of T cell epitope recognition by these memory T cells may provide more durable protection against emerging SARS-CoV-2 variants.

## Disclaimer

The authors declare no conflict of interest. Drs. E. Laing, M. Simons, C. Dalgard, A. Snow, R. Maves, D. Lindholm, C. Colombo, C. Lanteri, D. Tribble, T. Burgess and A. Malloy are service members or employees of the U.S. Government. This work was prepared as part of their official duties. Title 17 U.S.C. §105 provides that ‘Copyright protection under this title is not available for any work of the United States Government.’ Title 17 U.S.C. §101 defines a U.S.

Government work as a work prepared by a military service member or employee of the U.S. Government as part of that person’s official duties.

The contents of this publication are the sole responsibility of the author(s) and do not necessarily reflect the views, opinions, or policies of Uniformed Services University of the Health Sciences (USU); National Institutes of Health or the Department of Health and Human Services; the Henry M. Jackson Foundation for the Advancement of Military Medicine, Inc.; the Department of Defense (DoD); the U.S. Army Medical Department; the U.S. Army Office of the Surgeon General; the Departments of the Army, Navy, or Air Force; Brooke Army Medical Center; Walter Reed National Military Medical Center; Naval Medical Center San Diego; and Madigan Army Medical Center. Mention of trade names, commercial products, or organizations does not imply endorsement by the U.S. Government. The investigators have adhered to the policies for protection of human subjects as prescribed in 45 CFR 46.

## Acknowledgements

We thank Camille Estupigan for her editorial assistance in the preparation of this manuscript. We thank the members of the EPICC COVID-19 Cohort Study Group for their many contributions in conducting the study and ensuring effective protocol operations. The following members were all closely involved with the design, implementation, and oversight of the study.

Brooke Army Medical Center, Fort Sam Houston, TX: Col J. Cowden; S. Deleon; A. Markelz; K. Mende; T. Merritt; S. Merritt; R. Walter; CPT T. Wellington

Carl R. Darnall Army Medical Center, Fort Hood, TX: LTC S. Bazan; P. Kay Love

Fort Belvoir Community Hospital, Fort Belvoir, VA: L. Brandon; N. Dimascio-Johnson; MAJ E. Ewers; LCDR K. Gallagher; LCDR D. Larson; MAJ M. Odom; A. Rutt

Henry M. Jackson Foundation, Inc., Bethesda, MD: D. Clark

Infectious Disease Clinical Research Program, Uniformed Services University of the Health Sciences; Henry M. Jackson Foundation for the Advancement of Military Medicine, Inc., Bethesda, MD: G. Atwood; S. Banks; I. Barahona; L. Brandon; S. Cammarata; S. Chambers; S. Chi; S. Deleon; C. Fox; M. Grother; N. Kirkland; P. Kay Love; T. Lalani; K. Mende; T. Merritt; K. Miyasato; C. Morales; M. Oyeneyin; E. Parmelee; R. Resendez; W. Robb-McGrath; A. Rutt; M. Sanchez-Edwards; P. Sandoval; R. Tant

Madigan Army Medical Center, Joint Base Lewis McChord, WA: S. Chambers; CPT C. Conlon; CPT K. Everson; LTC P. Faestel; COL T. Ferguson; MAJ L. Gordon; LTC S. Grogan; CPT S. Lis; COL C. Mount; LTC D. Musfeldt; W. Robb-McGrath; MAJ R. Sainato; C. Schofield; COL C. Skinner; M. Stein; MAJ M. Switzer; MAJ M. Timlin; MAJ S. Wood

Naval Medical Center Portsmouth, Portsmouth, VA: G. Atwood; S. Banks; R. Carpenter; LCDR C. Eickhoff; CAPT K. Kronmann; T. Lalani; LCDR T. Lee; LCDR A. Smith; R. Tant; CDR T. Warkentien

Naval Medical Center San Diego, San Diego, CA: CAPT J. Arnold; CDR C. Berjohn; S. Cammarata; LCDR S. Husain; N. Kirkland; LCDR A. Lane; J. Parrish; G. Utz

Tripler Army Medical Center, Honolulu, HI: S. Chi; MAJ E. Filan; K. Fong; CPT T. Horseman; MAJ M. Jones; COL A. Kanis; LTC A. Kayatani; MAJ W. Londeree; LTC C. Madar; MAJ J. Masel; MAJ M. McMahon; K. Miyasato; G. Murphy; COL V. Ngauy; MAJ E. Schoenman; C. Uyehara; LTC R. Villacorta Lyew

Uniformed Services University of the Health Sciences, Bethesda, MD: C. Byrne; COL K. Chung;

C. Coles; D. Gunasekera; COL P. Hickey; LTC J. Livezey; COL T. Oliver; J. Rusiecki; A. Scher United States Air Force School of Medicine, Dayton, OH: A. Fries

Walter Reed National Military Medical Center, Bethesda, MD: I. Barahona; D. Gunasekera; M. Oyeneyin William Beaumont Army Medical Center, El Paso, TX: CPT M. Banda; CPT B. Davis; MAJ T. Hunter; CPT O. Ikpekpe-Magege; CPT S. Kemp; R. Mody; R. Resendez; P. Sandoval; COL M. Wiggins

