## Supplemental tables and figures for "Durability of SARS-CoV-2-specific T cell responses at 12-months post-infection"

### Table S1. Antibodies used for APC staining

| fluorochrome | Antigen | Vendor | Cat# |
|---|---|---|---|
| BUV395 | HLA-DR | BD biosciences | 564040 |
| L/D blue | L/D | | |
| BUV496 | CD56 | BD biosciences | 750479 |
| BUV563 | CD27 | BD biosciences | 748705 |
| BUV615 | CD206 | BD biosciences | 751638 |
| BUV661 | CD10 | BD biosciences | 741600 |
| BUV737 | CD69 | BD biosciences | 612817 |
| BUV805 | CXCR4 | BD biosciences | 742043 |
| BV421 | CD303 | BioLegend | 354212 |
| BV480 | Siglec 6 | BD biosciences | 747914 |
| BV510 | CD3 | BioLegend | 344828 |
| BV570 | CD11b | BioLegend | 301325 |
| BV605 | CD1c | BioLegend | 331538 |
| BV650 | CCR7 | BioLegend | 353234 |
| BV711 | CD25 | BioLegend | 356138 |
| BV750 | CD163 | BD biosciences | 747185 |
| BV785 | CD16 | BioLegend | 302046 |
| BB515 | CD141 | BioLegend | 565084 |
| PerCP | CXCR3 | BioLegend | 353740 |
| PerCP Cy5.5 | CD83 | BioLegend | 305320 |
| PE | CLEC9A | BioLegend | 353804 |
| PE-Dazzle 594 | CD11c | BioLegend | 337228 |
| PE-Cy5 | CD86 | BioLegend | 305408 |
| PE-Cy7 | CD80 | BioLegend | 375408 |
| AF700 | CD19 | BioLegend | 302226 |
| APC Cy7 | CD14 | BioLegend | 301820 |
| APC fire 810 | CD38 | BioLegend | 303550 |

### Table S2. The information of peptide pools used in the study

| Viral protein | Peptide pool name | Cat # | Vendor |
|---|---|---|---|
| SARS-CoV2-N | PepMix™ SARS-CoV-2 (NCAP) | PM-WCPV-NCAP | JPT |
| SARS-CoV2-VME | PepMix™ SARS-CoV-2 (VME1) | PM-WCPV-VME | JPT |
| SARS-CoV2-S | PepMix™ SARS-CoV-2 (Spike Glycoprotein SUB1) | PM-WCPV-S-SU1-1 | JPT |
| | PepMix™ SARS-CoV-2 (Spike Glycoprotein SUB2) | PM-WCPV-S-SU2-1 | JPT |
| SARS-CoV2-VEMP | PepMix™ SARS-CoV-2 (VEMP) | PM-WCPV-VEMP-1 | JPT |
| HKU1-S | PepMix™ HCoV- HKU1 (Spike Glycoprotein) | PM-HKU1-S | JPT |
| 229E-S | PepMix™ HCoV-229E (Spike Glycoprotein) | PM-229E-S | JPT |
| NL63-S | PepMix™ HCoV-NL63 (Spike Glycoprotein) | PM-NL63-S-1 | JPT |
| OC43-S | PepMix™ HCoV-OC43 (Spike Glycoprotein) | PM-OC43-S | JPT |
| CMV | CMV peptide pool for human CD4 and CD8 T cells | 3619-1 | Mabtech |

### Table S3. Antibodies used for T cell staining

| fluorochrome | Antigen | Vendor | Cat# |
|---|---|---|---|
| Surface | | | |
| BUV395 | HLA-DR | BD biosciences | 564040 |
| L/D blue | | ThermoFisher | L23105 |
| BUV563 | CD27 | BD biosciences | 748705 |
| BUV661 | PD1 | BD biosciences | 750260 |
| BUV737 | CD69 | BD biosciences | 612817 |
| BUV805 | CD28 | BD biosciences | 742037 |
| BV421 | CD3 | BD biosciences | 562877 |
| BV510 | CD62L | BioLegend | 304844 |
| BV650 | CCR7 | BioLegend | 353234 |
| BV750 | CXCR3 | BD biosciences | 746895 |
| BV785 | CD107a | BioLegend | 328644 |
| Spark Blue 550 | CD4 | BioLegend | 344656 |
| PerCP | CD8a | BioLegend | 301030 |
| PE Cy5 | CTLA4 | BD biosciences | 555854 |
| PE Cy5.5 | CD45RA | ThermoFisher | MHCD45RA18 |
| APC Fire 810 | CD38 | BioLegend | 303550 |
| Intracellular | | | |
| BUV615 | CD154 | BD biosciences | 751177 |
| BV480 | Ki67 | BD biosciences | 566109 |
| BV711 | IL13 | BD biosciences | 564288 |
| BB700 | FoxP3 | BD biosciences | 566526 |
| PE | IL21 | BioLegend | 513004 |
| PE Dazzle 594 | IL4 | BioLegend | 500832 |
| PE Cy7 | IFNγ | BioLegend | 506518 |
| APC | Granzyme B | ThermoFisher | MHGB05 |
| APC-R700 | IL17A | BD biosciences | 565163 |
| APC Fire750 | IL2 | BioLegend | 500352 |

### Table S4. COVID-19 study participant characteristics

| Code | Acute illness status | Sex | Race/ Ethnicity | Age (years) | Days hospitalized for COVID19 | Scale | Comorbidities | **Treatment**/ Routine medications/ Other | Vaccinated before collection |
|---|---|---|---|---|---|---|---|---|---|
| P01 | Inpatient | Male | Black | 52 | 5 | Conventional O2 | Obesity | | Yes |
| P03 | | Female | Black | 45.4 | 7 | | Borderline HTN | | No |
| P05 | | Male | Black | 47.5 | 4 | | Diabetes, asthma, obesity | ICS/LABA | No |
| P06 | | Male | White | 49.2 | 6 | | Chronic neurologic disorder | Ocrelizumab, InCS | No |
| P30 | | Male | White | 28.8 | 15 | | Leukemia, asthma, surgical asplenia | **Dexamethasone**, **remdesivir**, ICS | Yes |
| P17 | | Female | Black | 61.7 | 12 | High flow O2 | Chronic cardiac, pulmonary, and kidney disease, obesity | Dialysis | No |
| P20 | | Female | Asian | 52.7 | 8 | | Asthma | **convalescent plasma**, **prednisone**, ICS | Yes |
| P38 | | Male | Hispanic | 40.5 | 13 | | | **remdesivir** | No |
| P09 | | Female | Black | 56.2 | 13 | Invasive O2 | Graves disease, obesity | InCS | No |
| P41 | | Male | Asian | 21.4 | 4 | | Chronic hematologic disease | **Ionotropes, remdesivir**, renal replacement therapy, lisinopril | Yes |
| P02 | | Male | Black | 54.5 | 8 | No O2 | Obesity | | No |
| P04 | | Female | Black | 45.3 | 4 | | Obesity | Ibuprofen, Smoker | No |
| P07 | | Male | White | 57.6 | 2 | | HIV | Bictegravir/FTC/TAF, ICS | Yes |
| P08 | | Female | Black | 72.4 | 5 | | Diabetes | lisinopril | Yes |
| P11 | Outpatient | Female | White | 42 | | Minimal symptoms | Chronic neurologic disorder | | No |
| P14 | | Male | White | 54.4 | | | Cancer, HIV | valacyclovir, ABC/3TC/ dolutegravir, vape | Yes |
| P28 | | Male | White | 52.6 | | | Asthma | InCS | Yes |
| P32 | | Female | Hispanic | 40 | | | Obesity | | Yes |
| P13 | | Female | White | 56.3 | | Limited activity | Cancer | | Yes |
| P23 | | Female | Black | 46.4 | | | | | Yes |
| P25 | | Male | White | 60.2 | | | | | No |
| P33 | | Male | Asian | 36.2 | | | | | No |
| P34 | | Male | White | 39.3 | | | | | No |
| P35 | | Male | Native Hawaiian | 49.1 | | | | losartan | Yes |
| P36 | | Female | Hispanic | 40.3 | | | Diabetes | losartan | Yes |
| P37 | | Male | Hispanic | 32.9 | | | | | No |
| P39 | | Female | Multiple | 44.6 | | | | | Yes |
| P40 | | Female | Hispanic | 46 | | | Mild liver disease, cancer, obesity | | Yes |
| P42 | | Male | Hispanic | 20.1 | | | | | No |

Abbreviations:
ICS= inhaled corticosteroid,
LABA=long acting beta2-adrenergic agonist,
InCS=intranasal corticosteroids
HTN=hypertension
ACE=angiotensin converting enzyme
ABC=abacavir
3TC=lamivudine
FTC=emtricitabine
TAF=tenofovir alafenamide

Fig S1

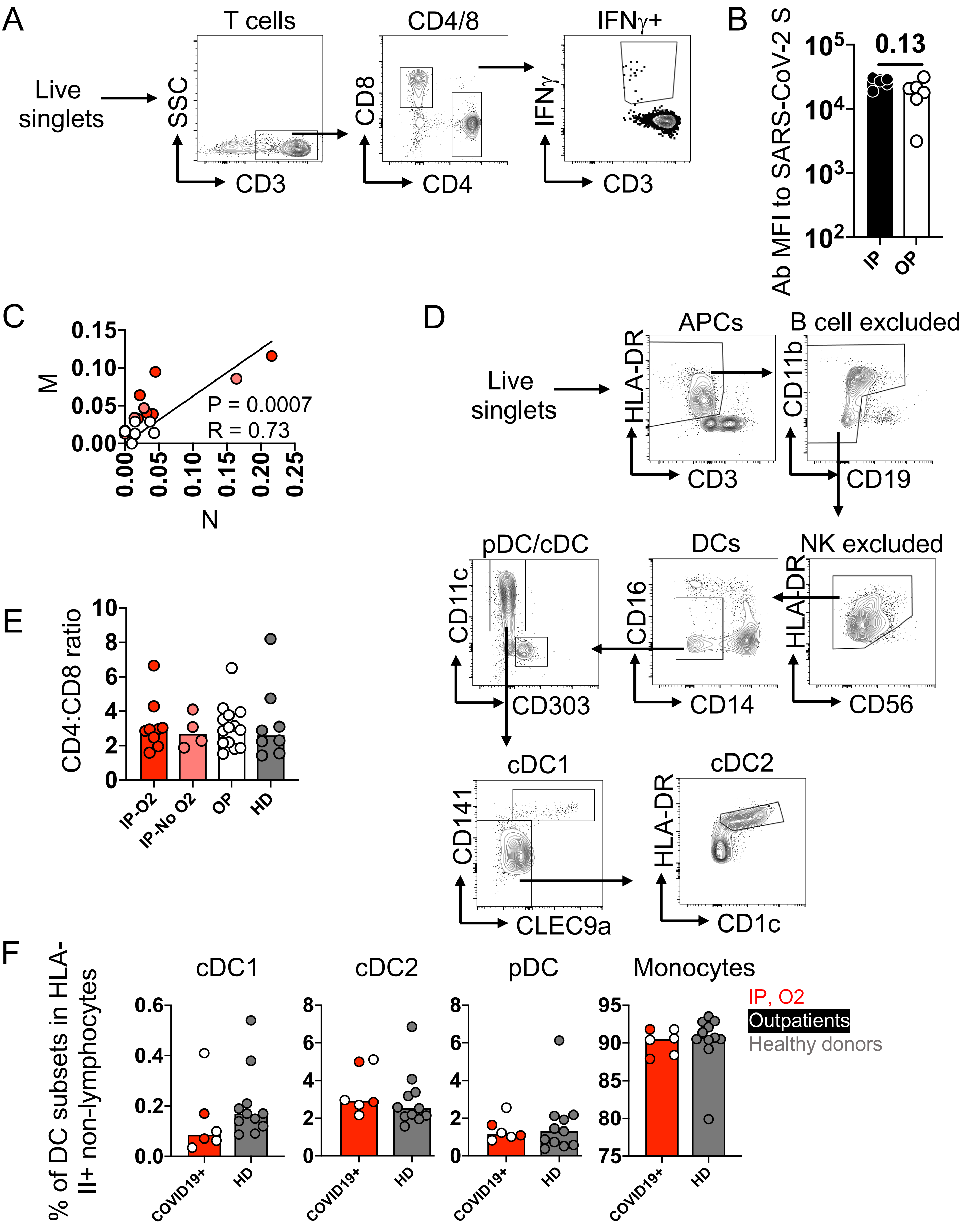
A
Live singlets
T cells
SSC
CD3
CD4/8
CD8
CD4
IFNγ+
IFNγ
CD3
B
Ab MFI to SARS-CoV-2 S
0.13
IP
OP
C
M
N
P = 0.0007
R = 0.73
D
Live singlets
APCs
HLA-DR
CD3
B cell excluded
CD11b
CD19
NK excluded
HLA-DR
CD56
DCs
CD16
CD14
pDC/cDC
CD11c
CD303
cDC1
CD141
CLEC9a
cDC2
HLA-DR
CD1c
E
CD4:CD8 ratio
IP-O2
IP-No O2
OP
HD
F
% of DC subsets in HLA-II+ non-lymphocytes
cDC1
cDC2
pDC
Monocytes
COVID19+
HD
IP, O2
Outpatients
Healthy donors

**Supplemental Figure 1. Investigation of antigen presenting cells (APCs) and major T cell subsets. A.** Gating strategy to characterize CD4 and CD8 T cells and antigen-specific T cells by expression of IFNγ. **B.** Antibodies to SARS-CoV-2 spike glycoprotein at 12-months post-infection. Participants vaccinated before blood collection were excluded. **C**. Correlation of T cell responses to SARS-CoV-2 N and M. P value in the plot indicates significance by Spearman rank correlation. R value in the plot indicates coefficient. **D.** Gating strategy to characterize the classical APCs in PBMCs. **E.** CD4:CD8 ratio among groups were similar. Red dots represent inpatients with oxygen supplementation, pink dots represent inpatients without oxygen supplementation, open dots represent outpatients, grey dots represent healthy donors, respectively. **F.** Frequencies of APC subsets including cDC1 and 2, plasmacytoid DCs, and monocytes in HLA-II+CD19- non-lymphocytes between COVID-19 patients and healthy donors.

Fig S2

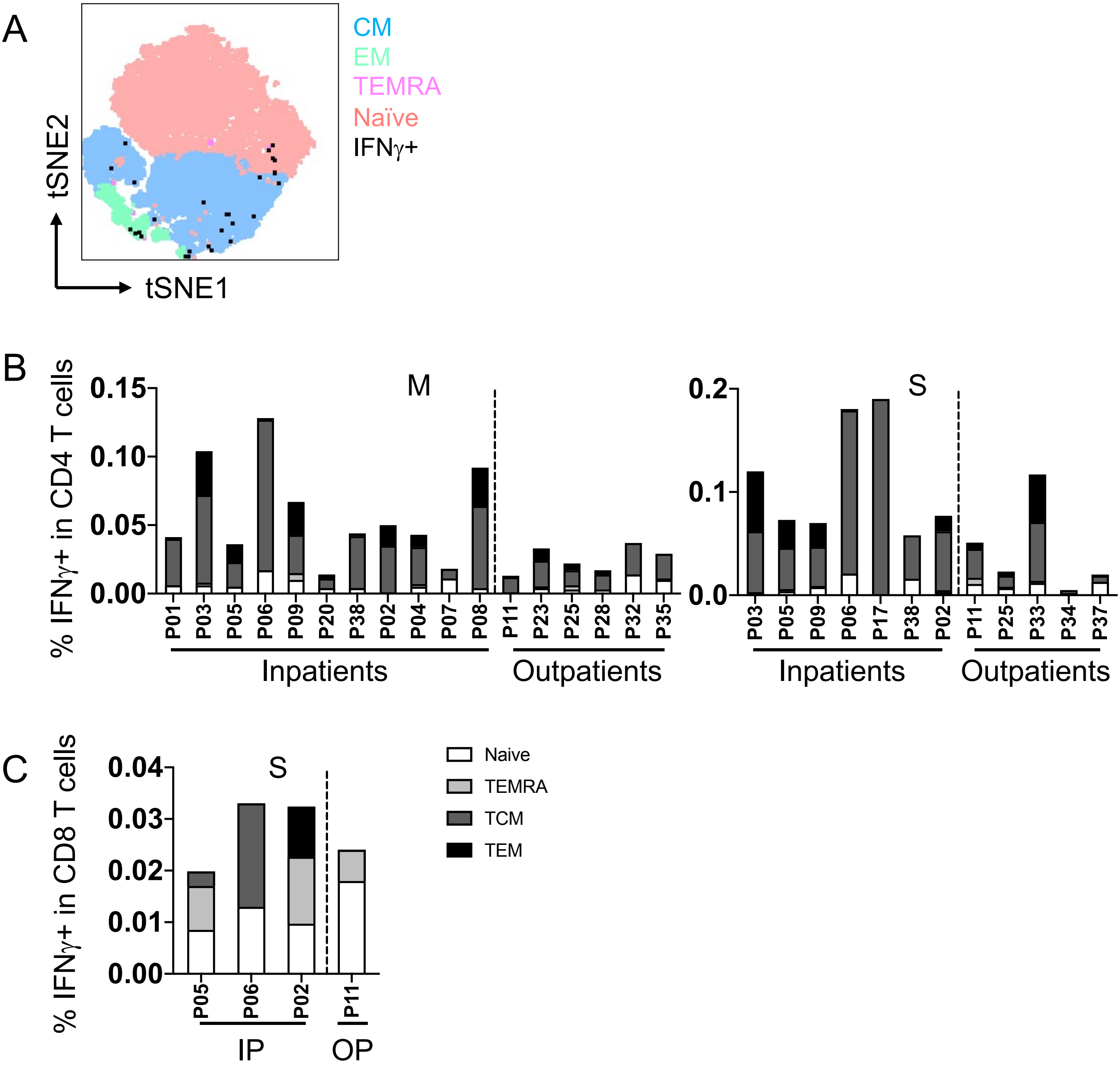


**Supplemental Figure 2. Memory phenotype characterization of SARS-CoV-2-specific T cell responses. A.** tSNE using memory markers CCR7, CD45RA, CD62L, CD27, and CD28 to show that IFNγ+ CD4 T cells were mainly CCR7+CD45RA-CD28+CD27+, indicating central memory phenotype. **B.** Similar to N-specific CD4 T cell memory phenotypes, M and S-specific CD4 T cells were mainly central memory. **C.** Similar to N-specific CD8 T cell memory phenotypes, S-specific CD8 T cells exhibited diverse memory phenotypes.

Fig S3

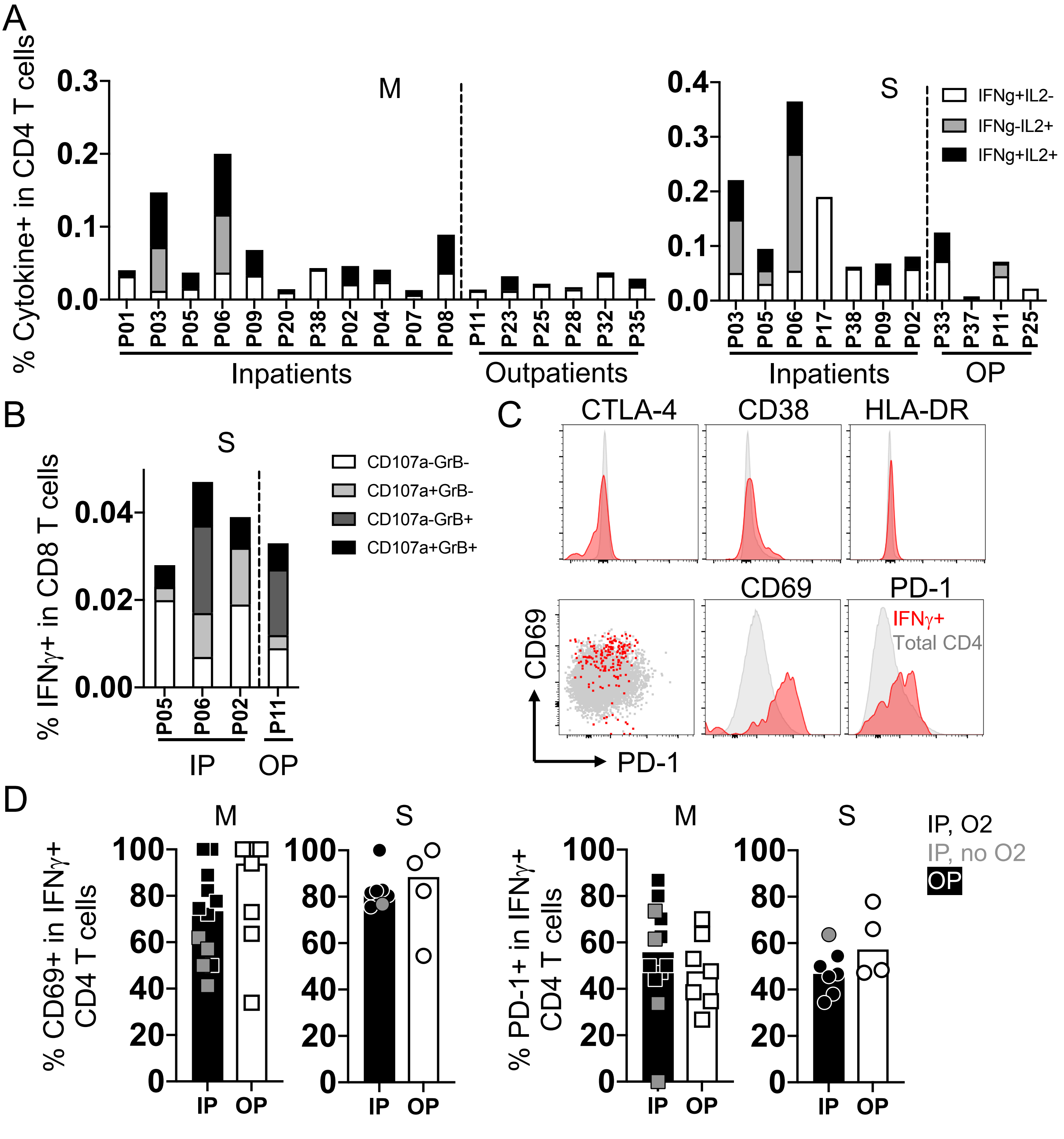


**Supplemental Figure 3. Polyfunctionality, cytotoxicity, and activation of SARS-CoV-2-specific CD4 or CD8 T cells. A.** Polyfunctionality of CD4 T cell responses to SARS-CoV-2 M and S proteins measured by IL2 and IFNγ. **B.** Cytotoxicity of IFNγ+ CD8 T cells to SARS-CoV-2 S protein measured by expression of intracellular Granzyme B and membrane CD107a. **C.** Histograms of the expression of T cell activation markers CTLA-4, CD38, HLA-DR, CD69, and PD-1, and a dot plot of expression of CD69 and PD-1 in parent and IFNγ+ CD4 T cells. Red indicates IFNγ+ CD4 T cells; grey indicates parent CD4 T cells. **D.** The frequencies of CD69 and PD-1 in SARS-CoV-2 M and S-specific IFNγ+ CD4 T cells in each patient group. Grey dots in the inpatient group represent inpatients without oxygen supplementation.
